# TLS_Finder: An algorithm for Identifying Tertiary Lymphoid Structures Using Immune Cell Spatial Coordinates

**DOI:** 10.1101/2024.12.26.630405

**Authors:** Ayse A Koksoy, Maria Esther Salvatierra, Patient Mosaic Team, Luisa Maren Solis Soto, Cara Haymaker

**Author notes:** **Corresponding Author:** Ayse A Koksoy, Department of Translational Molecular Pathology, The University of Texas, MD Anderson Cancer Center, 2130 West Holcombe Blv, Houston, TX, 77030. Collaborators: (Patient Mosaic Team) Nadim J Ajami, Azad Ali, Franklin Alvarez, Brittany Alvarez, Bianca Amador, Surosh Avandsalehi, Claudia Alvarez Bedoya, Elena Bogatenkova, Elizabeth Bonojo, Maria Neus Bota, Elizabeth M Burton, Noble Cadle, Vanessa Castro, Chi-Wan Chow, Randy Aaron Chu, Shadarra Crosby, Candace Cunningham, Carrie Daniel-MacDougall, Mary Domask, Sheila Duncan, Ellie Freebern, Andrew Futreal, Vivian Gabisi, Jessica Gallegos, Andrea Galvan, Jose Garcia, Ana Garcia, Celia Garcia-Prieto, Christopher Gibbons, Jonathan Benjamin Gill, Dominic Guajardo, Curtis Gumbs, Kristin J Hargraves, Cara Haymaker, Timothy Heffernan, Joshua Hein, Sharia Hernandez, Yasmine M Hoballah, Theresa Honey, Chacha Horombe, Cindy Hwang, MD (Habibul) Islam, Stacy Jackson, Jeena Jacob, Akshaya Jadhav, Robert Jenq, Weiguo Jian, Juliet Joy, Isha Khanduri, Walter Kinyua, Laura Klein, Mark Knafl, Larisa Kostousov, Diana Kouangoua, Ying-Wei Kuo, Wenhua Lang, Barrett Craig Lawson, Alexander Lazar, Jack Lee, Erma Levy, XiQi ‘Cece’ Li, Yang Li, Latasha D Little, Yang Liu, Yan Long, Vielka Lopez, Wei Lu, Sandra Lugo, Aaliyah Maldonado, Jared Malke, Asri Margono, Dipen Maheshbhai Maru, Grace Mathew, Surena Matin, Brian McKinley, Jennifer Leigh McQuade, Courtney McRuffin, Gertrude Mendoza, Francisco Montemayor, Raymond Montoya, Theresa T Nguyen, Heather Perez, Juan Posadas Ruiz, Sabitha Prabhakaran, Mallory Psenda, Gabriela Raso, Carl Rhodes, Mike Roth, Pranoti Sahasrobhojane, Amber Savant, Keri L Schadler, Alejandra Serrano, Kenna R Shaw, Julie M Simon, Elizabeth Sirmans, Luisa Maren Solis Soto, Xingzhi ‘Henry’ Song, Meghan Stennis, Huandong ‘Howard’ Sun, Maria Chang Swartz, Christopher Vellano, Angela Walker, Wanlin Wang, Ignacio Ivan Wistuba, Scott Eric Woodman, Xiaogang ‘Sean’ Wu, DeArtura ‘DeeDee’ Young, Jianhua ‘John’ Zhang, Haifeng Zhu, Hui ‘Helen’ Zhu. **Author Contributions:** Ayse A Koksoy – conceptualization, methodology, validation, writing original draft, visualization, supervision, project administration, writing (review and editing) Maria E Salvatierra - validation; writing (review and editing) Luisa Solis Soto - resources, writing (review and editing) Cara Haymaker - resources, writing (review and editing).

## Abstract

Tertiary lymphoid structures (TLS) are lymphoid formations that develop in non-lymphoid tissues during chronic inflammation, autoimmune diseases, and cancer. Accurate identification and quantification of TLS in tissue can provide crucial insights into the immune response of several disease processes including antitumor immune response. TLS are defined as aggregates of T cells, B cells and dendritic cells. In histological tissue sections stained with Hematoxylin and Eosin they are identified as aggregates of 50 or more lymphoid cells, however immunohistochemical analysis are required to confirm presence of distinct immune cell patterns. Assessment of lymphoid aggregates can be done in H&E slides or in slides stained with single or multiplex immunohistochemistry or other tissue based high-plex approaches with key biomarkers such as CD3 (T cells) and CD20 (B cells), however manual assessment of them is time consuming thus limiting its full evaluation. To our knowledge, published algorithms that identify TLS within a tissue are based on histological assessment of H&E slides or through use of ML algorithms trained on images that show the TLS presence in tissues; and quantification and spatial analysis of TLS still remains a challenge. This study aims to develop a robust algorithm to recognize TLS using the spatial coordinates of immune cells in any given tissue. The algorithm uses X and Y coordinates of T and B cells in tissues identified by a pathologist-supervised digital image analysis of multiplex chromogenic immunohistochemistry of tissues stained with CD3 and CD20. The algorithm is flexible to be used for detailed analysis of TLS stages; including other cell types within the definition of TLS; such as dendritic cells (DC) and high endothelial venules (HEV); to assess different stages of TLS formation.

## Introduction

Tertiary lymphoid structures (TLS) and lymphoid aggregates (LA) represent critical components of immune cell organization within the tumor microenvironment (TME). TLS are sites of lymphoid tissue generation in non-lymphoid tissues. Unlike primary and secondary lymphoid organs, TLS are formed de novo in tissues under pathological conditions such as chronic inflammation, autoimmune disorders and cancer (^1^,^2^) **Fig 1**. These structures are crucial for mounting effective adaptive immune responses, as they facilitate the local priming and activation of T cells and B cells, promoting antigen presentation and enhancing anti-tumor immunity. (^3^,^4^)

TLS have been characterized by distinct immune zones, including a germinal center supported by follicular dendritic cells (fDC) and high endothelial venules (HEV), which allow for the recruitment and retention of lymphocytes (^5^). While TLS exhibit variable compositions of immune cells and maturational stages; their formation is typically indicative of a robust local immune response against disease or tumor progression (^6^). Lymphoid aggregates are defined by presence of more than 50 T and/or B cells at a given spot and the TLS are more organized and defined by the presence of a high endothelial venule and follicular Dendritic cells (fDC) in the center of this bulk (^3,7^). The identification of TLS has historically relied on histological analysis, where fully matured structures are readily identifiable by pathologists in H&E-stained tissue slides(^8,9^). However, immature or less defined TLS, which lack the clear germinal centers or HEV, are more challenging to accurately detect and quantify(^10^). This challenge can be overcome by the development of computational tools capable of automating and standardizing TLS identification, thereby enhancing reproducibility and facilitating large-scale analyses.

Typically, the generation of adaptive immune response to chronic inflammatory diseases and cancer, involves infiltration of the tissue with lymphocytes and formation of new lymphoid structures (^11^). TLS presence in tumor microenvironment (TME) has been associated with transplant rejection, better pathogen clearance and higher anti-tumor response (^1^) . Presence of TLS and their simpler form, lymphoid aggregates; have been associated with better therapy response and superior patient outcomes in cancer(^12^). The stages and quantity of TLS, correlate with cancer stages and recurrence(^13^), their presence is especially valued in assessing prognosis of breast, colorectal and lung cancer(^14,15^). TLS have also been suggested to correlate and predict immune check point inhibitor efficacy in solid tumors (^16^) .Due to the prognostic and predictive potential of TLS; there has been an effort in the field to accurately quantitate their presence.

The current TLS annotating algorithms use machine learning and deep learning approach and their effectiveness can be limited by many factors including; the tissue type they are trained for finding TLS, details in labeling, size of the acquired images for training, artifacts in the images (^9,17,18^). The advent of spatially resolved immune profiling technologies, such as mIF and IHC has enabled researchers to study the intricate cellular architecture within TME with high resolution. In this context, spatial analysis algorithms like **TLS Finder**, provide an opportunity to move beyond subjective, labor-intensive histological methods. By leveraging immune cell phenotyping data in combination with their spatial coordinates, our algorithm efficiently identifies TLS based on the counts and proximity of T and B cells, capturing the key features of immune cell interactions within TME. Furthermore, the ability to incorporate other immune marker components such as the dendritic cells and HEV, enhances the algorithms’ flexibility, allowing it to detect fully mature TLS. This spatially driven approach aligns with emerging trends in cancer immunotherapy, where the spatial context of immune cell populations; rather than mere cell counts, has been shown to have predictive value for treatment response; particularly in an era of ICI therapies.

## Methods

The FFPE slides stained with multiplex IHC (3plex panel, Ventana) were used for marker staining. After annotation of images by the pathologist, a cell by marker dataframe with coordinates for each cell is formed. Marker concatenation per cell using Pandas (Python) defined cell phenotypes. ‘TLS Finder’ algorithm is a spatial analysis algorithm, designed to identify lymphoid structures (LS) such as lymphoid aggregates (LA) and TLS. The algorithm works by using the X and Y coordinates and the Python SciPy and Networkx modules, to measure presence of T and/or B cells within a circular area with a T or a B cell center.

The function takes a dataframe of cells with defined phenotypes or cell specific markers, cell coordinates (x,y) and a radius parameter (to define the neighborhood for spatial proximity analysis). KDTree is constructed using cell coordinates (x,y) to facilitate efficient querying. A graph ‘G’ is initialized where each cell represents a node. Edges are added between nodes of the graph if the specified nodes fall in the specified radius from a random seed. This step identifies clusters of TLS or LA of relevant cell types based on spatial proximity. The cells used in defining one cluster are not reused for a second time. The function outputs a dataframe with the identified aggregates, center coordinates and the corresponding TLS classification. The python code for the ‘TLS Finder’ algorithm can be found at https://github.com/AAKoksoy/TLS-Finder.

For this study, CD20 and CD8 markers were used to define B cells and T cells, respectively. The cell size was standardized to 10 microns, and a query radius of 160 microns was selected to account for possible empty spaces between cells. A lymphoid aggregate was defined as an area containing 50 or more B and/or T cells within a 160-micron radius of a central B or T cell. Cells already counted in one LA or TLS area were excluded from subsequent assessments. (Figure 1)

To assess the algorithm’s effectiveness in recognizing the circular structures with HEV in the center, we generated a synthetic dataset of 120000 cells (B-cell, T-cell, Other-cell, HEV), with one TLS at the center of the image comprised of one HEV at the center, surrounded by a circular placement of B-cell and an outer placement of T cells; representing the TLS Bulls-eye structure. The code for this data can also be found at our github page.

## Results

The algorithm was tested on 2 independent slides to assess if it is matching pathologist annotations for locating the TLS and used in 2 different studies to assess for presence of TLS (Figures 2 and 3). It successfully identifies immune cells and determines their spatial distributions. In 2 different test datasets; with different concentration and composition of cells in aggregate centers (Table 1); the algorithm identified the centers accurately, matching those of the pathologist annotations. With a query radius of 160 microns, 84 small lymphoid aggregate centers that were components of the 5 larger more complex structures were found, with the coordinates of each center provided. These cells and centers matched the findings from the pathologist annotations. These findings demonstrate the algorithm’s capability to detect and locate TLS within the tumor microenvironment.

**Figure 1.**
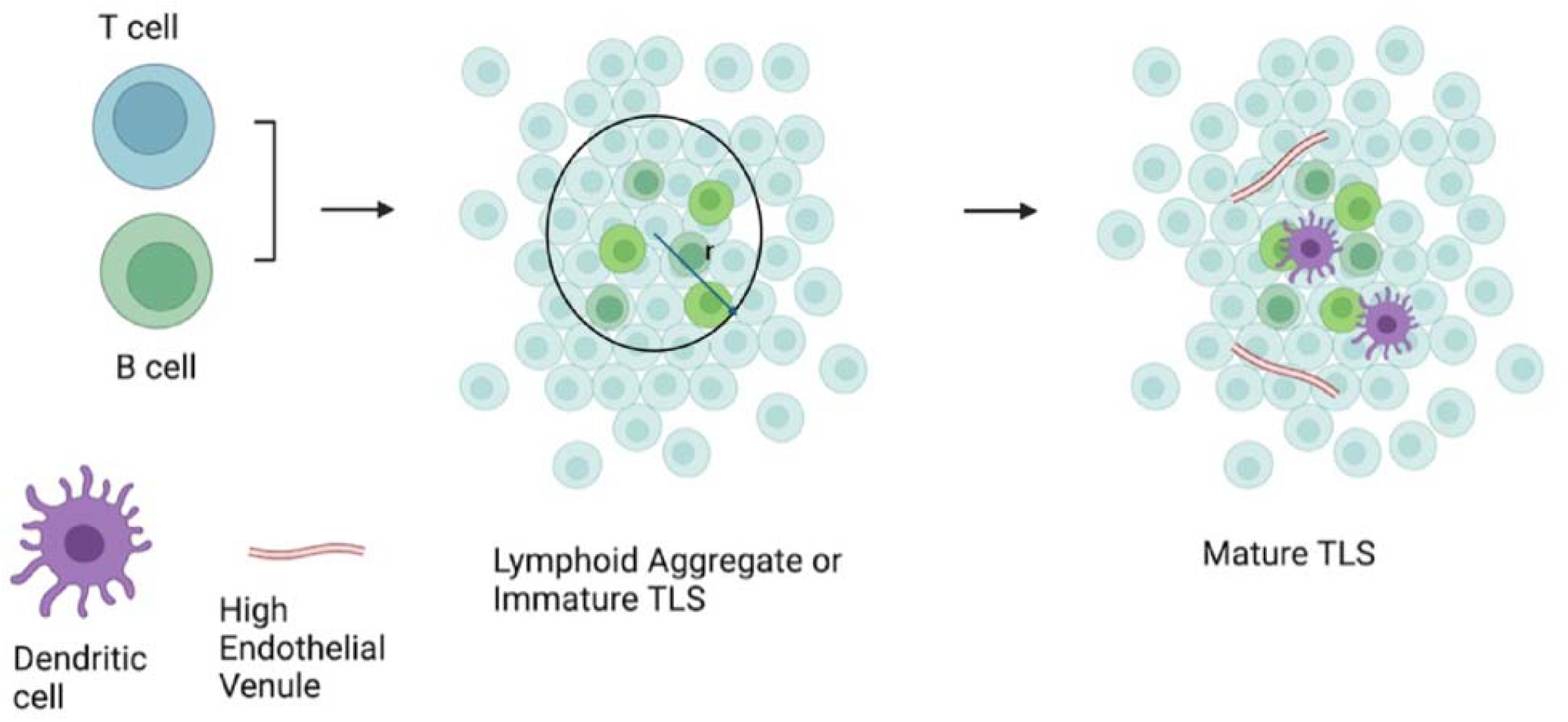
TLS schematic drawing. TLS are defined as presence of more than 50 T or B cells within a 60 micron radius from a T or B cell. Our algorithm used the cells that are CD8+ or CD20+ to assess the presence of TLS within a query radius of 60 microns as shown above.

**Figure 2.**
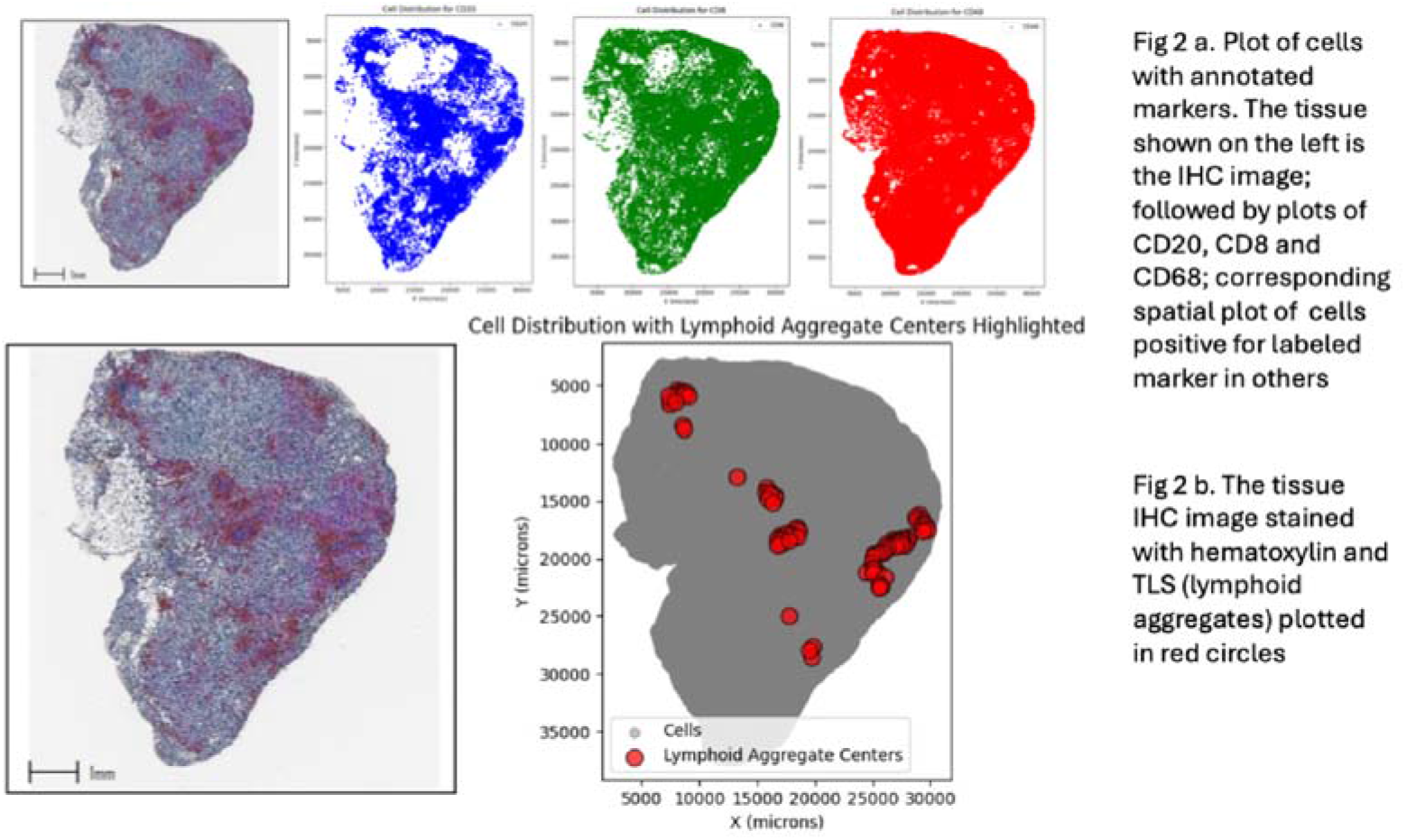
The original image and the cell types plotted for case 1.

**Figure 3.**
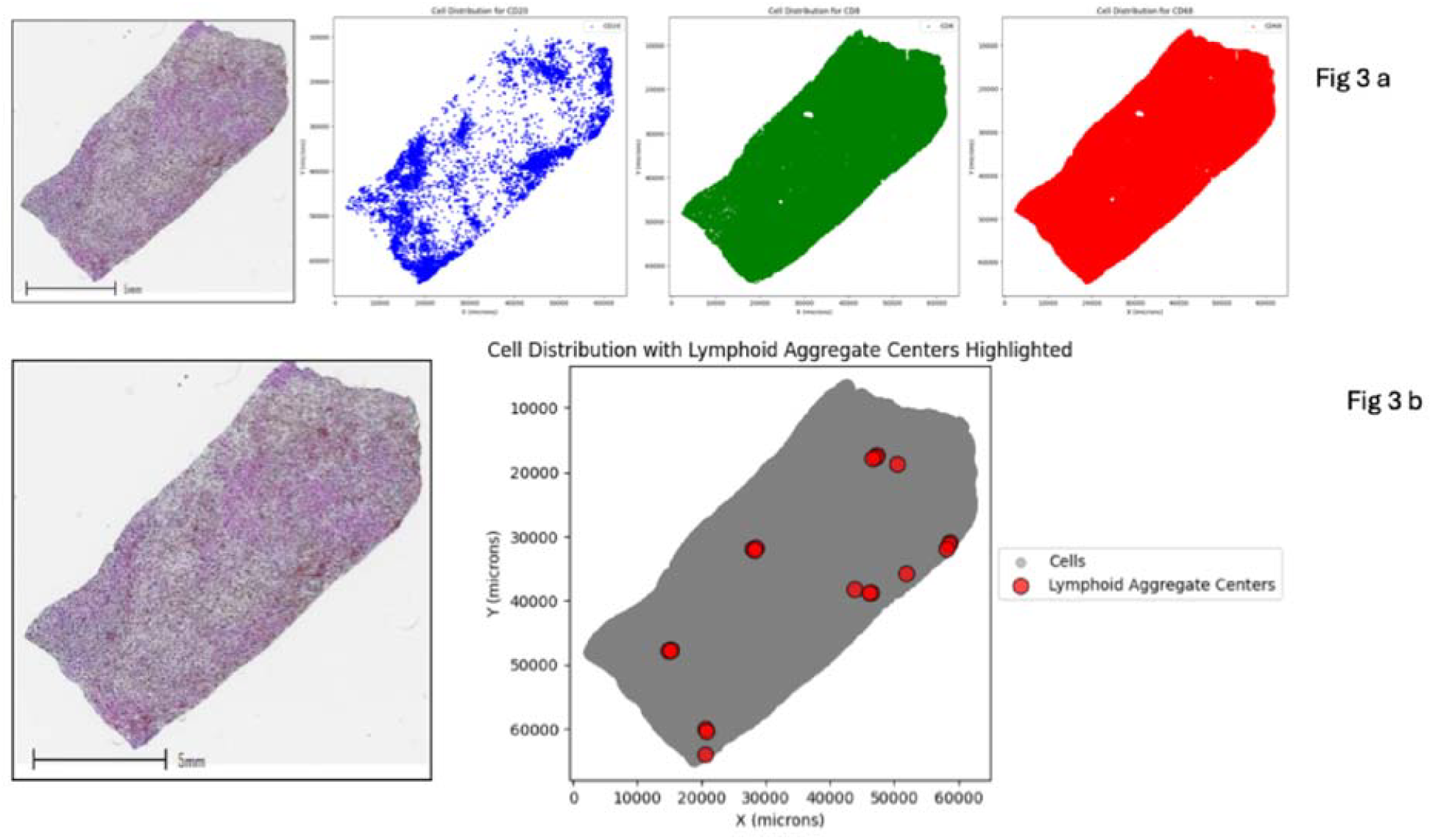
The original image and the cell types plotted for case 2.

**Table 1.**
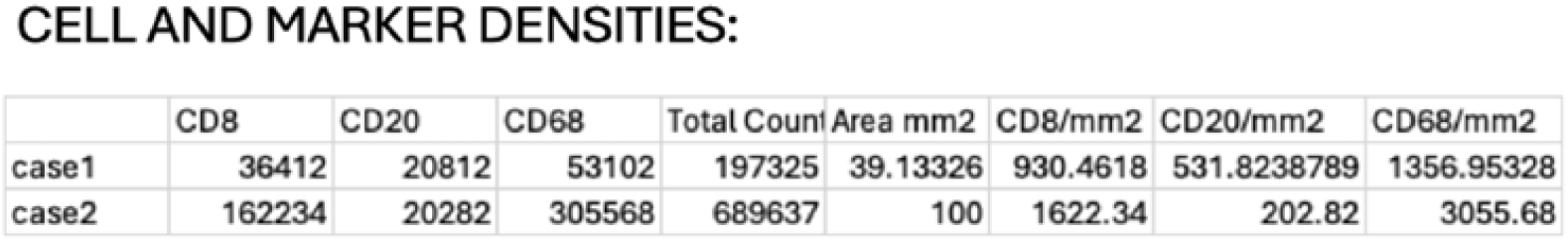
The marker densities raw and normalized to area.

## Supplementary Results and Figures

We also wanted to demonstrate the ability of the function to detect circular structures with specific cells in center such as mature TLS with and HEV at center. For this purpose we created a synthetic dataset using Python, Numpy module that complies with these descriptions. The synthetic data constructed a pseudo-tissue with a TLS in the center and random distribution of T cells, B cells and Other cells in the rest of the tissue. (Sup Fig1.) We were able to demonstrate that the algorithm captures the location of the TLS correctly and also locates the Lymphoid aggregates present in the pseudo-tissue data.

## Discussion

The algorithm developed and tested in this study provides an effective tool for identifying lymphoid structures based on the spatial distribution of CD8+ and/or CD20+ immune cells. This approach leverages the spatial proximity of immune cells to identify the TLS, by-passing the need for subjective and time-consuming annotation process using H&E slides. Importantly, the algorithm is flexible and can incorporate additional markers, such as dendritic cells, and high endothelial venules (HEV) and follicular dendritic cells (fDC) to show fully mature TLS, if necessary. Our results show that the TLS Finder algorithm accurately matches pathologist annotations, providing confidence in its reliability to detect lymphoid aggregates. The 160 micron radius used in this study effectively captures the spatial organization of LA in TME. Given the prognostic and predictive potential of TLS in various cancers, including breast, colorectal, and lung; this capability of defining LA and TLS by using cell biomarker positivity and spatial coordinates becomes particularly relevant.

By utilizing spatial proximity data to detect immune cell clusters, TLS Finder brings several advantages over traditional methods and overcomes their limitations. This approach enables the identification of both fully mature and immature TLS, which may lack the characteristic features such as germinal centers. Previous studies have highlighted challenges in identifying TLS solely based on histological appearance or immunohistochemistry; particularly when these structures are not fully formed to their mature state. Considering most of the pathology image data are of immunohistochemistry (IHC) or immunofluorescence (IF) origin; algorithm’s ability to analyze presence of TLS from X and Y coordinates of cells in image data of IHC and multiplex immunofluorescence (mIF) to generate precise location and coordinates of immune clusters overcomes these challenges and provides another landmark to further analyze spatial interactions within TME. Moreover, the algorithm is able to handle large datasets generated from high-throughput and/or WSI of IHC or mIF platforms, which makes it an ideal component for studying complex immune landscape of tumors. Algorithm is robust so that data from translational platforms comprised of phenotyped cells with X and Y coordinates can also be used. The only limitation of the algorithm is that it requires correct phenotyping of cells prior.

Given the established association between TLS and improved patient outcomes, as well as their potential role in predicting immune checkpoint inhibitors; the TLS Finder algorithm represents a valuable tool for translational cancer research. We are presenting the algorithm with the hope that it may be helpful in refining the prognostic models and therapeutic decision-making; and ultimately contribute to more personalized treatment strategies.

## Supplemental Data and Figures

**Supplementary Fig 1.**
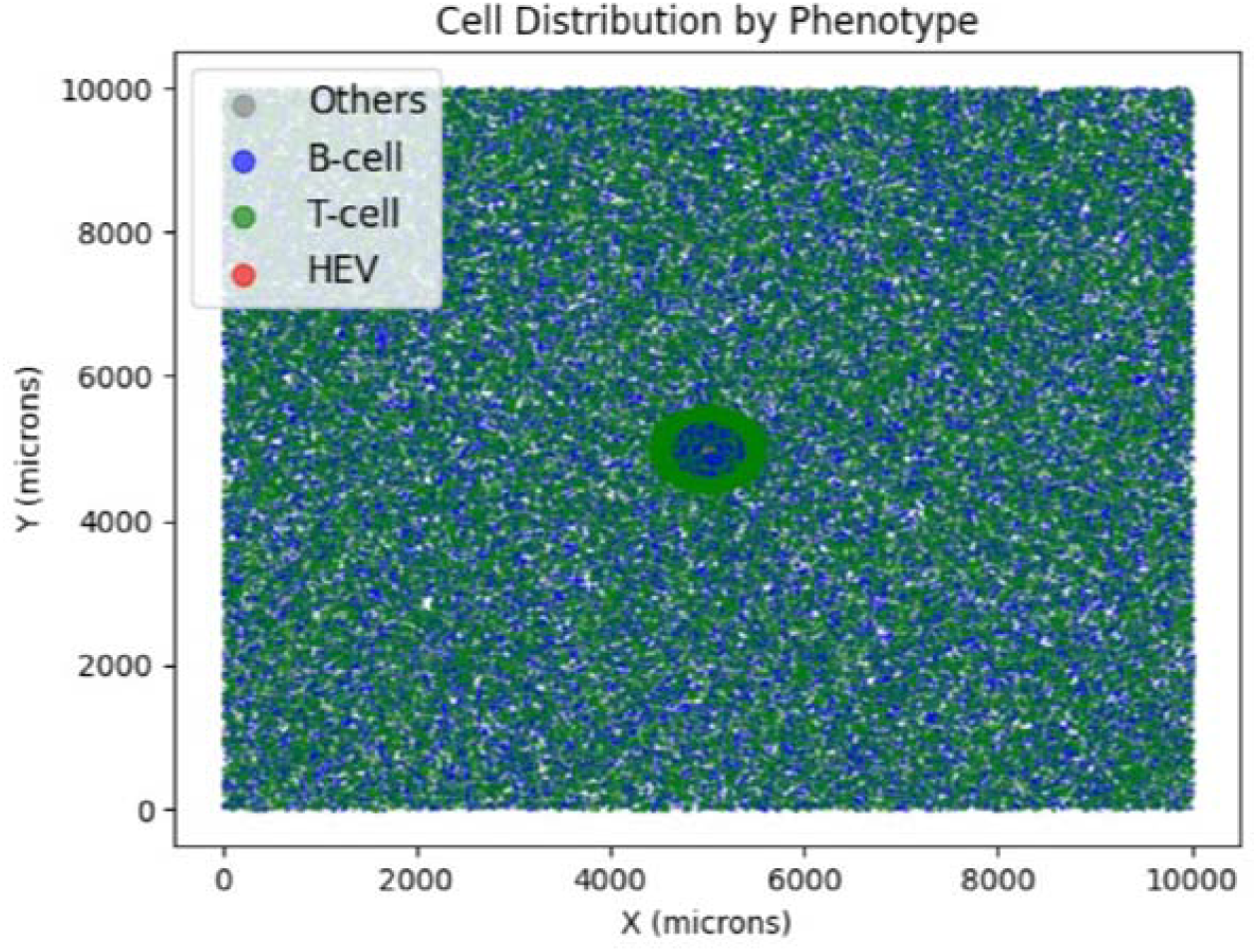
Pseudo-tissue image created by synthetic dataset. The mature TLS with HEV shown by the cyan at the center of the pseudo-tissue image.

**Supplementary Fig 2.**
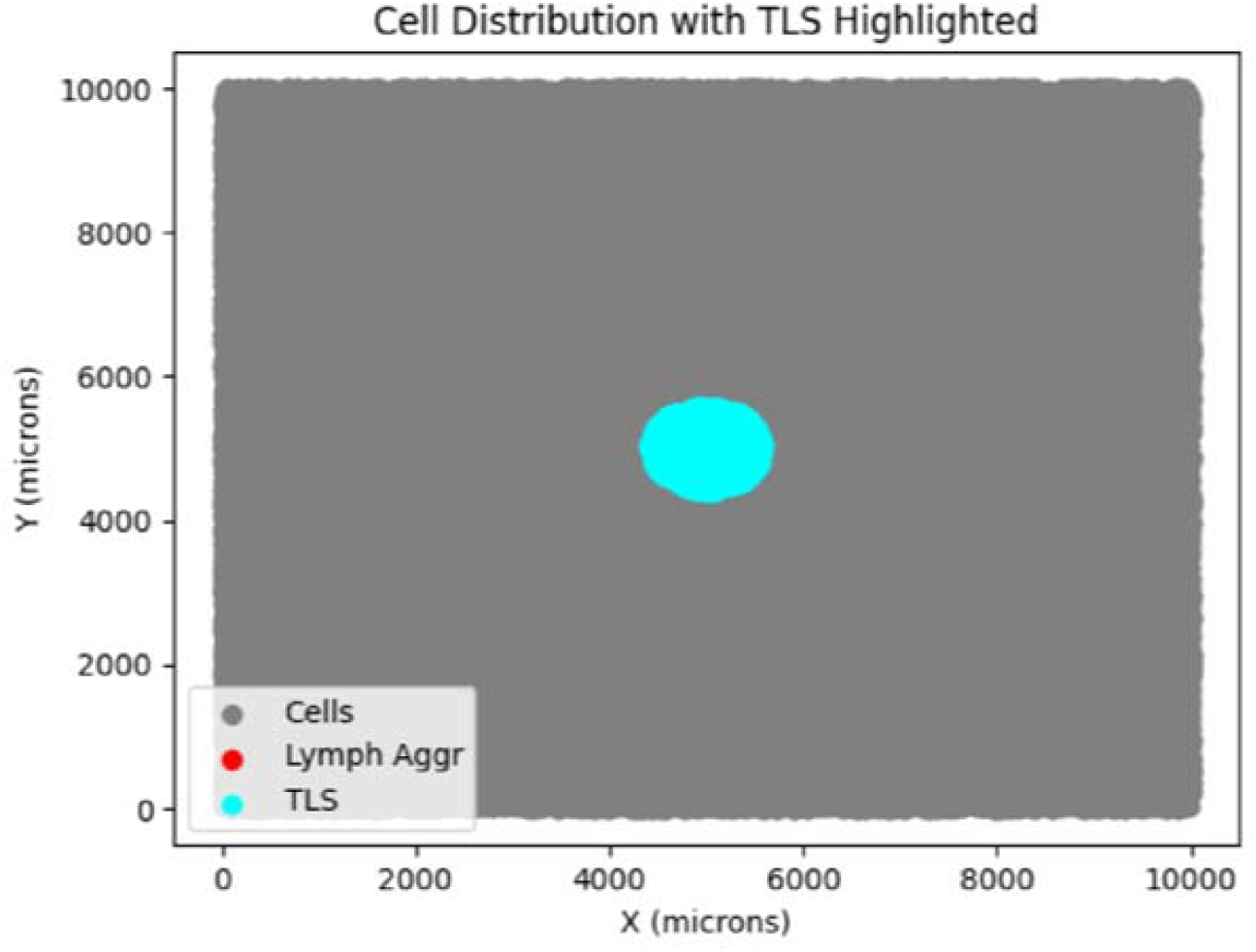
The mature TLS marked by the cyan at the center of the pseudo-tissue image.

**Supplementary Fig 3.**
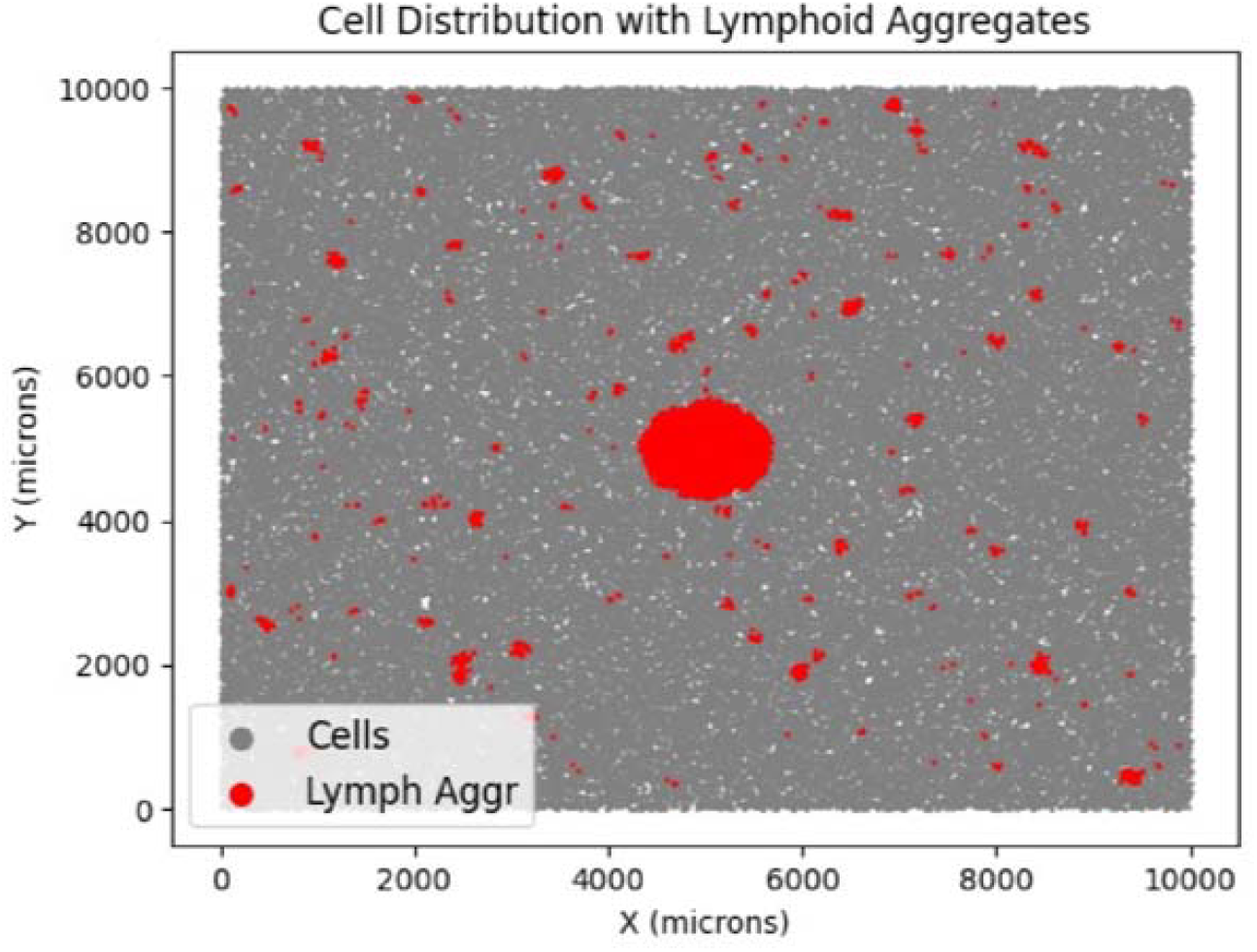
The ‘Lymphoid Aggregates’ marked by red throughout the pseudo-tissue image.

## Notes

**Funding:** This project was supported in part by the NIH CCSG Award (CA016672 (Institutional Tissue Bank (ITB) and Research Histology Core Laboratory (RHCL)), the Translational Molecular Pathology-Immunoprofiling lab (TMP-IL) MoonShots Platform at the Department Translational Molecular Pathology, the University of Texas MD Anderson Cancer Center.

https://github.com/AAKoksoy/TLS-Finder

### Competing Interest Statement

The authors have declared no competing interest.

### Summary of Updates

Patient Mosaic Team name was corrected.

## References

1. Dieu-Nosjean, M. et al. Tertiary lymphoid structures, drivers of the anti-tumor responses in human cancers. Immunological Reviews 271, 260–275 (2016).

2. Sato, Y., Silina, K., Van Den Broek, M., Hirahara, K. & Yanagita, M. The roles of tertiary lymphoid structures in chronic diseases. Nat Rev Nephrol 19, 525–537 (2023).

3. Schumacher, T. N. & Thommen, D. S. Tertiary lymphoid structures in cancer. Science 375, eabf9419 (2022).

4. Sautès-Fridman, C., Petitprez, F., Calderaro, J. & Fridman, W. H. Tertiary lymphoid structures in the era of cancer immunotherapy. Nat Rev Cancer 19, 307–325 (2019).

5. Pipi, E. et al. Tertiary Lymphoid Structures: Autoimmunity Goes Local. Front. Immunol. 9, 1952 (2018).

6. Helmink, B. A. et al. B cells and tertiary lymphoid structures promote immunotherapy response. Nature 577, 549–555 (2020).

7. Sue-Chu, M. et al. Lymphoid Aggregates in Endobronchial Biopsies from Young Elite Cross-country Skiers. Am J Respir Crit Care Med 158, 597–601 (1998).

8. Lin, J.-R. et al. High-plex immunofluorescence imaging and traditional histology of the same tissue section for discovering image-based biomarkers. Nat Cancer 4, 1036–1052 (2023).

9. Barmpoutis, P. et al. Tertiary lymphoid structures (TLS) identification and density assessment on H&E-stained digital slides of lung cancer. PLoS ONE 16, e0256907 (2021).

10. Vanhersecke, L. et al. Standardized Pathology Screening of Mature Tertiary Lymphoid Structures in Cancers. Laboratory Investigation 103, 100063 (2023).

11. Munoz-Erazo, L., Rhodes, J. L., Marion, V. C. & Kemp, R. A. Tertiary lymphoid structures in cancer – considerations for patient prognosis. Cell Mol Immunol 17, 570–575 (2020).

12. Lynch, K. T. et al. Heterogeneity in tertiary lymphoid structure B-cells correlates with patient survival in metastatic melanoma. J Immunother Cancer 9, e002273 (2021).

13. Posch, F. et al. Maturation of tertiary lymphoid structures and recurrence of stage II and III colorectal cancer. OncoImmunology 7, e1378844 (2018).

14. Song, I. H. et al. Predictive Value of Tertiary Lymphoid Structures Assessed by High Endothelial Venule Counts in the Neoadjuvant Setting of Triple-Negative Breast Cancer. Cancer Res Treat 49, 399–407 (2017).

15. Rakaee, M. et al. Tertiary lymphoid structure score: a promising approach to refine the TNM staging in resected non-small cell lung cancer. Br J Cancer 124, 1680–1689 (2021).

16. Vanhersecke, L. et al. Mature tertiary lymphoid structures predict immune checkpoint inhibitor efficacy in solid tumors independently of PD-L1 expression. Nat Cancer 2, 794–802 (2021).

17. Komura, D. & Ishikawa, S. Machine Learning Methods for Histopathological Image Analysis. Computational and Structural Biotechnology Journal 16, 34–42 (2018).

18. Van Rijthoven, M. et al. Multi-resolution deep learning characterizes tertiary lymphoid structures and their prognostic relevance in solid tumors. Commun Med 4, 5 (2024).

